# Polyclonal sensory neuron derivation from iPSCs as an efficient alternative to single clone strategies for pain-relevant *in vitro* models

**DOI:** 10.64898/2026.09.08.750017

**Authors:** Anika Neureiter, Luise Rüßmann, Petra Hautvast, Maike F. Dohrn, Noortje W. M. van den Braak, Arzu Stoppe, Martin Häusler, Annette Lischka, Katja Eggermann, Angelika Lampert

## Abstract

Induced pluripotent stem cell (iPSC) workflows typically rely on monoclonal cultures, requiring multiple independent cell lines to compensate for clonal variability. This increases workload, cost, and limits scalability for translational applications. Here, we establish polyclonal reprogramming strategies using Sendai virus to generate clonally diverse iPSC cultures already during reprogramming through FACS, MACS, manual selection, or prolonged culture. Both monoclonal and FACS-derived polyclonal cultures reached pluripotency. Reprogramming transgenes were silenced, whereas Sendai virus (SeV) mRNA persisted across passages in both culture types; heat treatment at 38.5 °C markedly reduced SeV levels. Monoclonal and polyclonal cultures differentiated efficiently into neural crest-like cells and sensory neurons. As a second polyclonal strategy, monoclonal cultures were pooled at day-5 of sensory neuron differentiation to generate percentage-controlled progenitor mixtures. TRPA1 protein expression has not been shown reliably in iPS-derived sensory neurons. We hypothesized that increased clonal diversity might facilitate detection of TRPA1 protein expression. Despite rigorous antibody validation, TRPA1 protein expression was detectable only diffusely across the whole cell area. Together, our results show that polyclonal iPSC strategies enable clonally inclusive generation of patient-specific sensory neurons. These workflows reduce early workload and provide a robust foundation for scalable drug-screening and *patient-in-a-dish* applications.

## 1. Introduction

The discovery of induced pluripotent stem cells (iPSCs) in 2006 and 2007 ^1,2^ fundamentally transformed biomedical research by enabling the generation of patient-specific human cell models. Since iPSCs retain the donor’s genetic and epigenetic background while maintaining the capacity to differentiate into virtually any human somatic cell type, they provide a powerful platform for studying individual disease mechanisms and for developing personalized therapeutic approaches ^3^.

However, iPSC-based models are challenged by substantial biological and technical variability. Donor dependent differences and those introduced during reprogramming, clonal selection, long-term culture, and lineage specification can lead to pronounced heterogeneity between iPSC lines. This variability affects differentiation efficiency, cellular identity, and functional properties, thereby limiting reproducibility and complicating the interpretation of disease-related phenotypes ^4–9^.

Monoclonal iPSC strategies have been established as a promising approach to reduce this heterogeneity ^10^. Individual clones from the same donor still differ markedly in genetic stability, epigenetic state and functional characteristics not only on a clonal but also on single cell level ^7,11^. As a result, relying on a single clone per donor is insufficient to draw robust biological conclusions. Multiple independently derived iPSC lines are required to capture intrinsic variability and to ensure that observed phenotypes reflect real disease mechanisms rather than clonal artifacts ^12^.

Generating and characterizing several monoclonal lines per donor is, however, highly labor-, time-, and cost-intensive. This poses a substantial barrier in translational and preclinical research settings, where high throughput and time- and cost-efficient iPSC generation is essential ^3^. The challenge becomes even more pronounced when disease modelling requires highly specialized cell types such as sensory neurons, the differentiation of which from fully reprogrammed iPSCs adds an additional ∼10 weeks to the overall experimental timeline and further delays downstream applications such as drug screenings ^6,13–17^.

Polyclonal reprogramming strategies offer a practical way to overcome the limitations of monoclonal workflows by omitting the labor-intensive-step of single clone isolation, expansion, quality control and differentiation ^18,19^. In polyclonal cultures, fully reprogrammed iPSCs were reported to outgrow somatic cells and partially reprogrammed intermediates, allowing the establishment of pluripotent lines without the need to manually select individual colonies ^18^. Alternatively, automated selection of pluripotent cells from freshly reprogrammed polyclonal cultures have been published ^19^. Both strategies would enable the inclusion of clonal diversity in the experimental workflow, thereby maintaining biological representativeness by creating a *patient-in-a-dish* with far lower resource demands than manually selecting multiple clonal cell lines per donor.

Here, we aim to evaluate polyclonal reprogramming and differentiation strategies as an alternative to conventional monoclonal workflows for future modelling of neuropathic pain. To this end, we reprogrammed peripheral blood mononuclear cells (PMCS) from two chronic pain patients using both the standard monoclonal approach and a polyclonal reprogramming strategy. We assessed whether monoclonal and polyclonal iPSC lines successfully acquired pluripotency, maintained genetic integrity, and cleared the Sendai virus and reprogramming transgenes within a reasonable time frame. We further examined whether both approaches support efficient differentiation into sensory neurons. During differentiation, we additionally implemented a polyclonal strategy in which monoclonal- derived sensory neuron progenitors were mixed in defined proportions to preserve clonal diversity.

To challenge our polyclonal system, we focused on expression of TRPA1, a chemosensitive and mechanosensitive ion channel that plays a key role in nociception by detecting noxious stimuli and contributing to inflammatory and neuropathic pain signalling ^20^. Since TRPA1 expression has so far been difficult to achieve reliably in iPSC-derived sensory neurons ^21,22^, we selected TRPA1 as a test case to examine whether rare protein expression events are clone dependent. If only a subset of iPSC clones is capable of inducing TRPA1 surface expression during sensory neuron differentiation, monoclonal cultures may fail to capture this phenotype, whereas polyclonal or mixed-progenitor approaches could preserve TRPA1-competent clones within the population. By selecting TRPA1 as a model for rare and potentially clone-restricted protein expression, this study aims to determine whether polyclonal reprogramming and differentiation strategies can overcome monoclonal culture linked workload associated limitations and, additionally, better capture biologically relevant phenotypes than traditional monoclonal workflows.

## 2. Results

### 2.1 Reprogramming of monoclonal and polyclonal iPSCs

To systematically compare monoclonal and polyclonal reprogramming workflows, we established a unified experimental pipeline (Figure 1A) in which PBMCs were reprogrammed using Sendai virus vectors delivering OCT4, SOX2, KLF4 and MYC and subsequently processed through different isolation strategies. As illustrated in Panel A, manually picked monoclonal colonies served as the conventional reference, while the remaining reprogramming dish was used to generate polyclonal cultures that were either left unselected, manually selected or enriched for pluripotent cells using magnetic activated cell sorting (MACS, Miltenyi Biotech) or fluorescent activated cell sorting (FACS). This setup allowed us to directly assess how each isolation method influences the purity, morphology, and stability of emerging iPSC populations.

**Figure 1.**
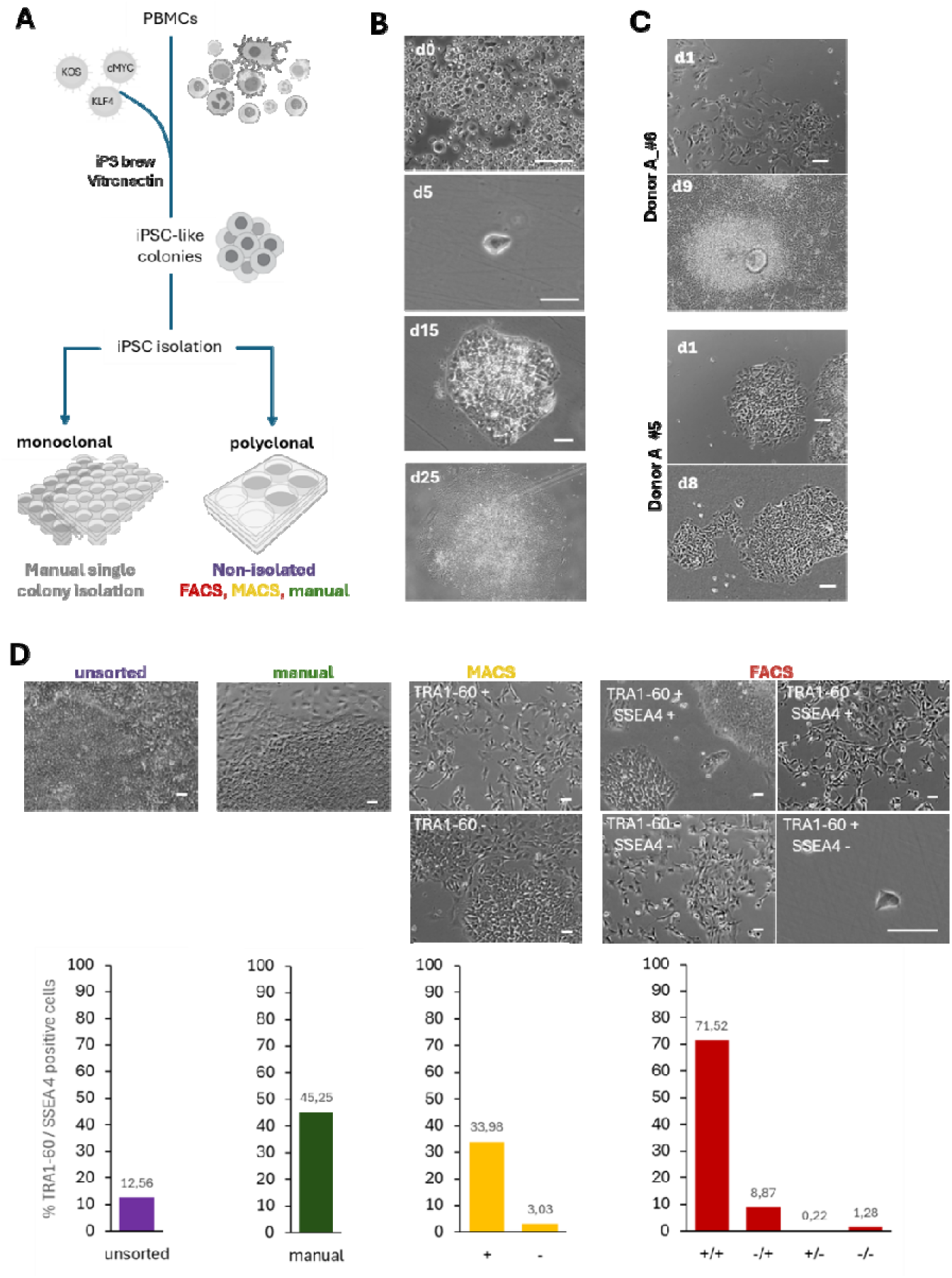
Reprogramming and purification of polyclonal induced pluripotent stem cells during reprogramming. **(A)** Schematic overview of monoclonal and polyclonal reprogramming workflows and isolation strategies for successfully reprogrammed cells. Images were taken from Bioicons (iPSCs icon by almaCELL is licensed under CC-BY 4.0 Unported https://creativecommons.org/licenses/by/4.0/) or NIH BioArt (Virus: NIAID Visual & Medical Arts. (5/22/2026). Arenavirus Silhouette. NIAID NIH BIOART Source. bioart.niaid.nih.gov/bioart/681, Cells: NIAID Visual & Medical Arts. (10/7/2024). Immune Cells. NIAID NIH BIOART Source. bioart.niaid.nih.gov/bioart/253 and 24- Wellplate: NIAID Visual & Medical Arts. (10/7/2024). 24 Well Plate. NIAID NIH BIOART Source. bioart.niaid.nih.gov/bioart/4. 6-Wellplate: NIAID Visual & Medical Arts. (10/7/2024). 6 Well Plate. NIAID NIH BIOART Source. bioart.niaid.nih.gov/bioart/6.) **(B)** Following Sendai virus transduction (d0), PBMCs progressively acquire iPSC-like morphology until day 15. **(C)** Clonally selected iPSCs one day and eight days after manual transition to clonal culture. DonorA_(#6) shows excessive expansion of differentiated cells causing exclusion of the clone from further experiments. DonorA_(#5) one day and 8 days after clonal selection showing typical iPS-morpholgy. After manual isolation of single iPSC colonies, remaining colonies were subjected to fluorescence-activated cell sorting (FACS) (Supplementary Figure 1). **(D**) One week after isolation, only FACS-purified TRA-1-60⁺/SSEA4⁺ cells display typical iPSC morphology, whereas MACS-isolated and unsorted cultures contain heterogeneous cell types. Quantification of TRA-1-60/SSEA4 expression by FACS one week after isolation demonstrates that only FACS-sorted iPSCs maintain high purity. Scale bar represents 100µm.

After infection of PBMCs with the reprogramming viruses, sporadic attachment of single cells was observed around day 5 (Figure 1B), followed by the emergence of compact iPSC-like colonies by day 15. Since each colony is assumed to originate from a single reprogrammed PBMC, the initial reprogramming dish represents a polyclonal mixture of independently reprogrammed cells. From this dish, 20 colonies resembling successfully reprogrammed cells were manually collected and expanded (exemplary morphology see Figure 1C). 14 of those successfully reprogrammed monoclonal cultures established morphologically a pluripotent state over the next weeks and maintained morphology throughout expansion (Figure 1C, lower panel). Remaining six clones were excluded due to presence of contaminating cells (Figure 1C, upper panel). Six presumptive pluripotent clones were selected for further quality control (Figure 2, Figure 3).

**Figure 2.**
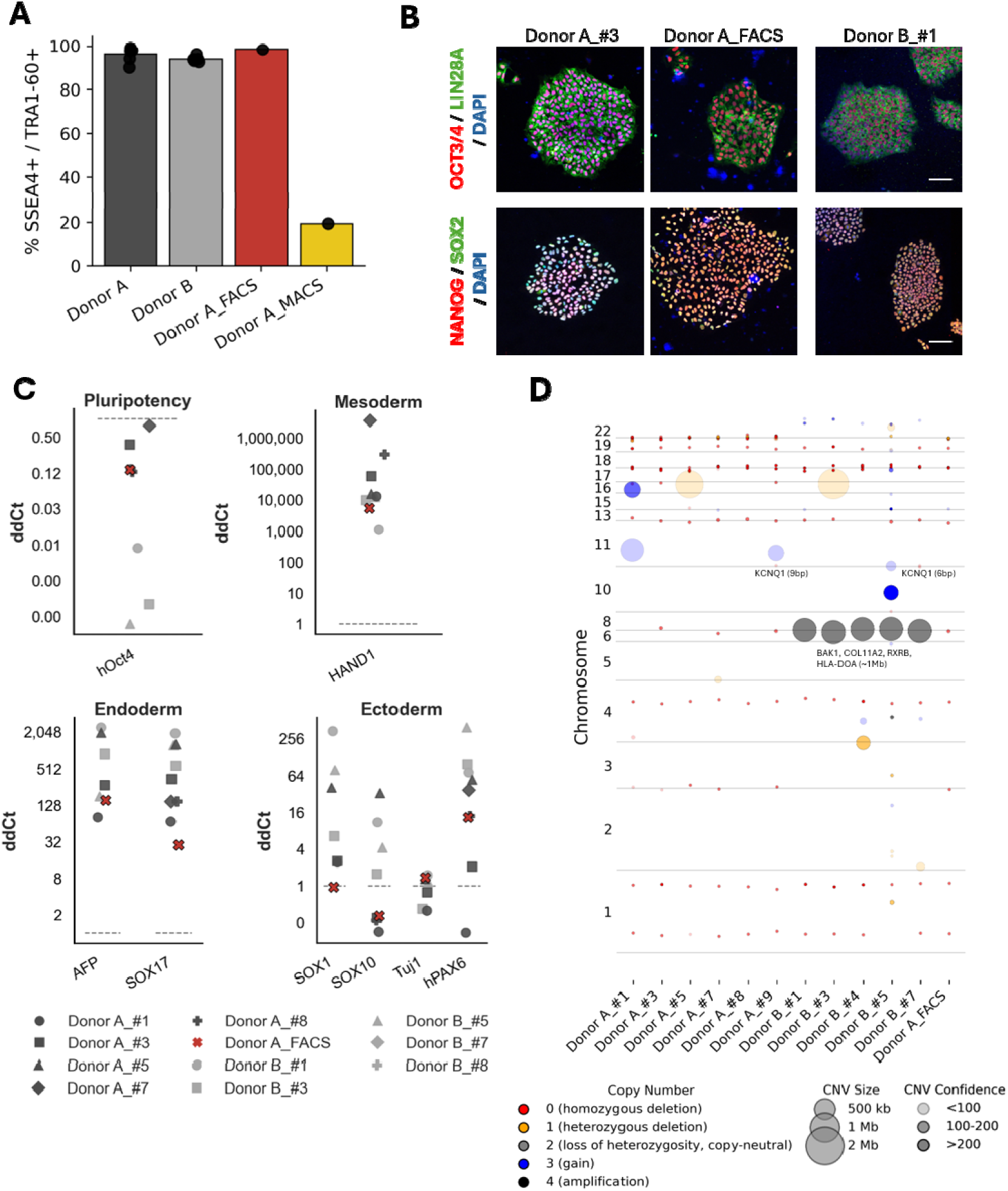
polyclonal and monoclonal iPSC-lines acquire pluripotency. Due to lower reprogramming efficiency, only monoclonal iPSC lines could be established from Donor B, while Donor A underwent polyclonal and monoclonal reprogramming. **(A)** All monoclonal iPSC lines from both donors expressed the pluripotency associated surface markers TRA-1-60 and SSEA4 as determined by flow cytometry. The FACS-derived polyclonal culture from Donor A showed marker expression comparable to monoclonal lines, whereas the MACS-derived polyclonal culture displayed poor marker expression and was excluded from further experiments. Background fluorescence was set to 0.5% based on isotype controls for each cell line. Percentages of TRA-1-60⁺/SSEA4⁺ double-positive cells are shown. **(B)** Immunofluorescence staining confirmed expression of the pluripotency markers OCT4, SOX2, NANOG and LIN28A in monoclonal iPSC lines and in the FACS-derived polyclonal culture. Scale bar: 100Dµm. Stainings for all monoclonal lines are provided in the Supplementary Information. **(C)** After 14 days of directed differentiation into the three germ layers, both monoclonal and polyclonal iPSC lines upregulated lineage-specific markers of ectoderm, mesoderm and endoderm, while downregulating OCT4. ΔΔCt values were normalized to the housekeeping genes GAPDH and HPRT and calculated relative to day-0 samples (dottet line). **(D)** Copy number variation (CNV) analysis of all iPSC lines revealed the genomic location (y-axis), size (dot size), copy number state and confidence of detected CNVs for each sample

**Figure 3.**
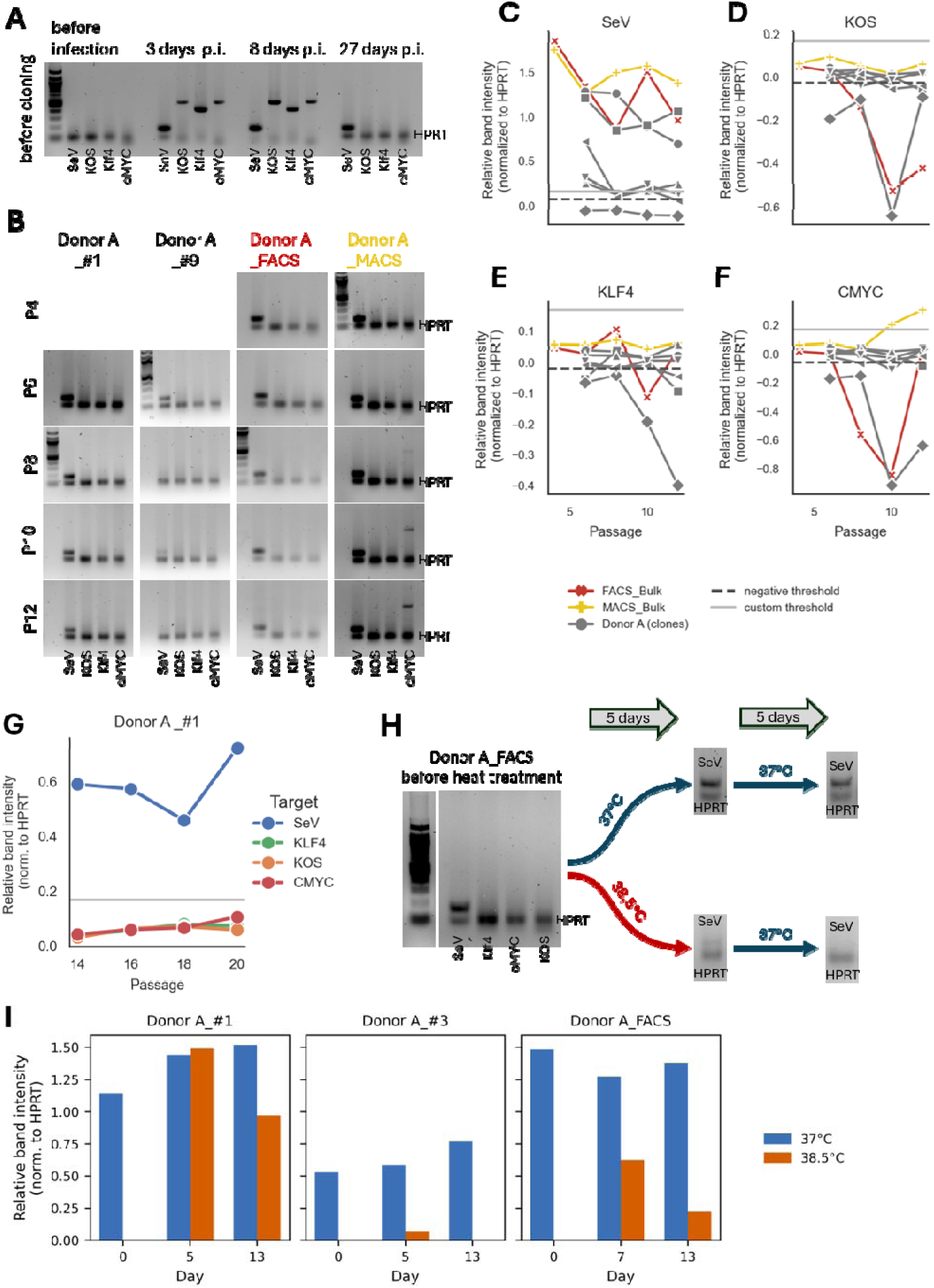
Sendai Virus and Trangene Clearance in monoclonal and polyclonal iPSC-cultures from Donor. **A.** (A) PCR followed by agarose gel electrophoresis was used to detect Sendai virus (SeV) and reprogramming transgenes (KLF4, cMYC, KLF4/OCT4/SOX2 (KOS) during the reprogramming process. HPRT (102-bp) served as loading control. Analysis of PBMCs before and 3, 8 and 27 days after Sendai virus transduction showed that only the SeV amplicon (181-bp) remained detectable prior to iPSC isolation. (B) After clonal isolation or MACS- and FACS-based enrichment, SeV expression persisted in some monoclonal iPSC lines, while others showed complete loss of SeV. Both FACS- and MACS-derived polyclonal cultures retained SeV expression. Notably, MACS-derived cultures also showed upregulation of cMYC (532-bp). (C–F) PCR bands were quantified, background-corrected and normalized to HPRT. Relative expression values are shown. KOS, cMYC and KLF4 remained below the optical detection threshold (light blue line) and within the range of negative controls across all conditions. In contrast, SeV expression remained high in both polyclonal cultures and in two monoclonal lines up to passage 12. (G) Extended passaging of a representative monoclonal line (Donor A #1) up to passage 20 did not result in SeV clearance. (H–I) FACS_Bulk and the monoclonal lines Donor A #1 and Donor A #3 were exposed to 38.5D°C for 5 days followed by recovery at 37 °C. Heat treatment resulted in reduced SeV expression in all three lines. Uncropped agarose gels are available in Supplementary Figure 3 and 4.

To establish polyclonal cultures, remaining cells after manual picking were expanded until 34 days after transduction and subjected to magnetic activated cell sorting (MACS, Miltenyi Biotech) based on TRA-1-60 expression or fluorescent activated cell sorting (FACS) based on TRA-1 60 and SSEA4 expression. Marker-expressing and negative populations were collected. FACS additionally allowed isolation of SSEA4-only populations, whereas TRA-1-60⁺/SSEA4⁻ cells were rarely detected (Figure 1D, Supplementary Figure 1).

One week after isolation, only FACS-enriched cultures showed absence of contaminating cells and formed densely packed colonies with sharp borders (Figure 1D), whereas MACS-isolated, manually collected and unselected cultures showed heterogeneous cell populations (Figure 1D). To determine if cells maintained marker expression, one week after isolation, the percentage of TRA-1-60 and SSEA4 expressing cells per selection method were assessed by flow cytometry (Figure 1D). MACS-purified cultures contained substantial numbers of contaminating cells and showed low percentages of TRA-1-60/SSEA4 double positive cells (34%). Manually collected polyclonal cultures displayed similarly modest purity (∼45%). In contrast, FACS sorted TRA-1-60⁺/SSEA4⁺ cells maintained high purity, with ∼70% of cells retaining expression of both markers and only ∼1% remaining double-negative (Figure 1D).

Together, these data demonstrate that FACS-based enrichment is, in our hands, the only method that reliably establishes cell cultures with high percentages of pluripotency-associated marker expression and colony morphology in polyclonal cultures soon after isolation, making it the most robust strategy for establishing high-quality *patient*-in-a-*dish* iPSC populations.

### 2.2 Monoclonal and FACS derived polyclonal iPSC-lines exhibit robust pluripotency and genetic stability

The *patient-in-a-dish* polyclonal approach is heavily influenced by reprogramming efficiency. The reprogramming efficiency differed between donors. Here, we reprogrammed cells from two donors, a father (Donor A) and his daughter (Donor B), both affected by a familial chronic pain syndrome characterized by widespread pain and pain hypersensitivity without clinical evidence of small- or large-fiber neuropathy. All attempts to reprogram cells of donor B resulted in low numbers of iPS-like colonies before isolation (Figure 2A, C, Supplementary Figure 2A,C);therefore, only monoclonal iPSC lines were established for this donor to allow further studies.

All monoclonal and the polyclonal MACS- and FACS- derived cell lines were subjected to flow cytometry after three to four weeks after initial isolation. All monoclonal lines from both donors expressed the pluripotency-associated surface markers TRA-1-60 and SSEA4 (Figure 2A). The FACS-derived polyclonal culture showed comparable levels of double-positive cells, whereas the MACS-derived polyclonal culture displayed markedly reduced marker expression with only 20% of cells co-expressing both markers. This polyclonal strategy was, therefore, excluded from further experiments, due to lack of efficiency.

Immunofluorescence staining demonstrated that monoclonal iPSC lines and the FACS-derived polyclonal culture expressed the core pluripotency factors OCT4, SOX2, NANOG and LIN28A (Figure 2B, Supplementary Figure 2B,C). All transcription factors (OCT4, SOX2 and NANOG) showed typical nuclear localization patterns while the translational enhancer LIN28A showed cytoplasmic localization. All cell lines exhibited compact colony morphology consistent with a pluripotent state.

To assess functional competence, monoclonal and polyclonal iPSC lines were subjected to three-germ-layer differentiation for 14 days. All lines downregulated OCT4 and upregulated lineage-specific markers of ectoderm, mesoderm and endoderm, confirming their trilineage potential (Figure 2C). ΔΔCt values normalized to GAPDH and HPRT and calculated relative to day 0 samples demonstrated comparable differentiation potentials across donors and monoclonal and polyclonal iPSC-cultures.

Genomic integrity was assessed by copy number variation (CNV) analysis (Figure 2D). All iPSC lines displayed largely stable genomes, with most CNVs occurring in both donors and across multiple cell lines. Donor A carried a recurrent ∼1 DMb CNV on chromosome 6 present in all derived lines. This CNV affected several genes, including BAK1, COL11A2, RXRB, and HLA-DOA, and was classified as a copy-neutral loss of heterozygosity. In addition, two monoclonal lines from Donor A exhibited a small 6–9Dbp gain in KCNQ1, a voltage-gated potassium channel predominantly expressed in cardiac tissue and epithelia. The FACS-derived polyclonal line closely resembled the monoclonal lines of Donor A in its CNV profile. Notably, unlike three of the sic monoclonal lines, the polyclonal culture did not carry any large CNVs, suggesting that polyclonal reprogramming preserves patient-specific CNVs comparably to monoclonal approaches, while clonal selection may favour the establishment of lines originating from clones with larger genomic alterations due to a selection bias.

Together, these data demonstrate that FACS-derived polyclonal iPSCs exhibit pluripotency marker expression, differentiation potential and genomic stability comparable to monoclonal iPSC lines, supporting their suitability for downstream applications.

### 2.3 SeV and transgene clearance in monoclonal and polyclonal iPSCs

Reprogramming of PBMCs was performed using a Sendai viral (SeV) vector system expressing the four Yamanaka factors OCT4, SOX2, KLF4, and cMYC. Transient expression of these factors is sufficient to induce pluripotency, but persistent expression of SeV or transgenes can interfere with the iPS-cell stage and downstream differentiation ^23,24^. Therefore, efficient elimination of these factors is essential for generating high-quality iPSC lines.

To monitor SeV and transgene clearance, RNA samples were collected throughout the reprogramming time course and analysed by PCR following the manufacturer’s recommendations. HPRT served as loading control. As expected, SeV and all four transgenes were absent before reprogramming and became detectable shortly after infection (day-3 post infection, p.i.; Figure 3A). By day-27 p.i., only SeV remained detectable, while all transgenes had already been eliminated.

To obtain SeV-free monoclonal lines, at least two rounds of subcloning are typically recommended. In our dataset, 2/6 monoclonal lines retained SeV also after passage 12, 2/6 lost SeV between passages 6 and 8, and 2/6 maintained SeV expression up to passage 12. Both FACS- and MACS-derived polyclonal cultures also showed residual SeV at all analyzed time points (Figure 3B). Quantification of PCR bands revealed that the amount of residual SeV relative to HPRT varied between lines. While SeV levels decreased in one monoclonal line, the other monoclonal line and both polyclonal cultures reached their lowest SeV expression around passage-6 (Figure 3C–F). All transgenes remained below the custom detection threshold in all lines except for the MACS-derived polyclonal culture, in which cMYC expression re-emerged at passage-8 and continued to increase.

Since the SeV clearance could require longer culture time, we extended the culture period of one representative monoclonal line (Donor-A_#1) to passage-20. SeV expression remained stable throughout all passages, whereas all transgenes stayed below the detection threshold (Figure 3G).

The SeV vectors used in this system contain temperature-sensitive mutations designed to facilitate viral clearance at 38–39 °C. To test this, SeV positive lines (Donor-A #1, Donor-A #3 and FACS_Bulk) were cultured at 38.5 °C for 5-days, followed by recovery at 37°C. In all lines, SeV levels decreased over time under heat treatment, whereas cultures maintained at 37°C showed stable SeV expression (Figure 3H–I). Donor-A_#3 showed complete loss of SeV after 5-days at 38.5 °C, and the FACS derived polyclonal culture reduced SeV to ∼20% of cells at 37°C. Donor-A_#1 retained SeV but at a markedly reduced level. These findings indicate that short-term culture at 38.5 °C promotes SeV elimination in both monoclonal and polyclonal iPSC cultures and should be considered as standard procedure during Sendai virus mediated reprogramming.

### 2.4 Differentiation of monoclonal and polyclonal iPSC cultures into sensory neurons

Polyclonal iPSC cultures are particularly attractive for *patient-in-a-dish* disease modelling and drug screening, as they preserve patient-specific cellular diversity; however, such applications require robust differentiation into the relevant somatic cell type. As the patients involved in this study are pain patients, we differentiated both monoclonal and polyclonal iPSC lines into sensory neurons.

To assess whether polyclonal iPSC cultures can be differentiated into sensory neurons comparable to monoclonal iPSC-cultures, iPSCs from both donors were first induced to form neural crest-like cells (NCLCs; Figure 4A) ^21,25^. For this, iPSCs were cultured as free-floating spheres in the presence of EGF and FGF2. After several days in suspension, spheres attached to uncoated plastic surfaces and NCLCs migrated outward from all iPSC lines, indicating comparable neural crest induction across monoclonal and polyclonal cultures.

**Figure 4.**
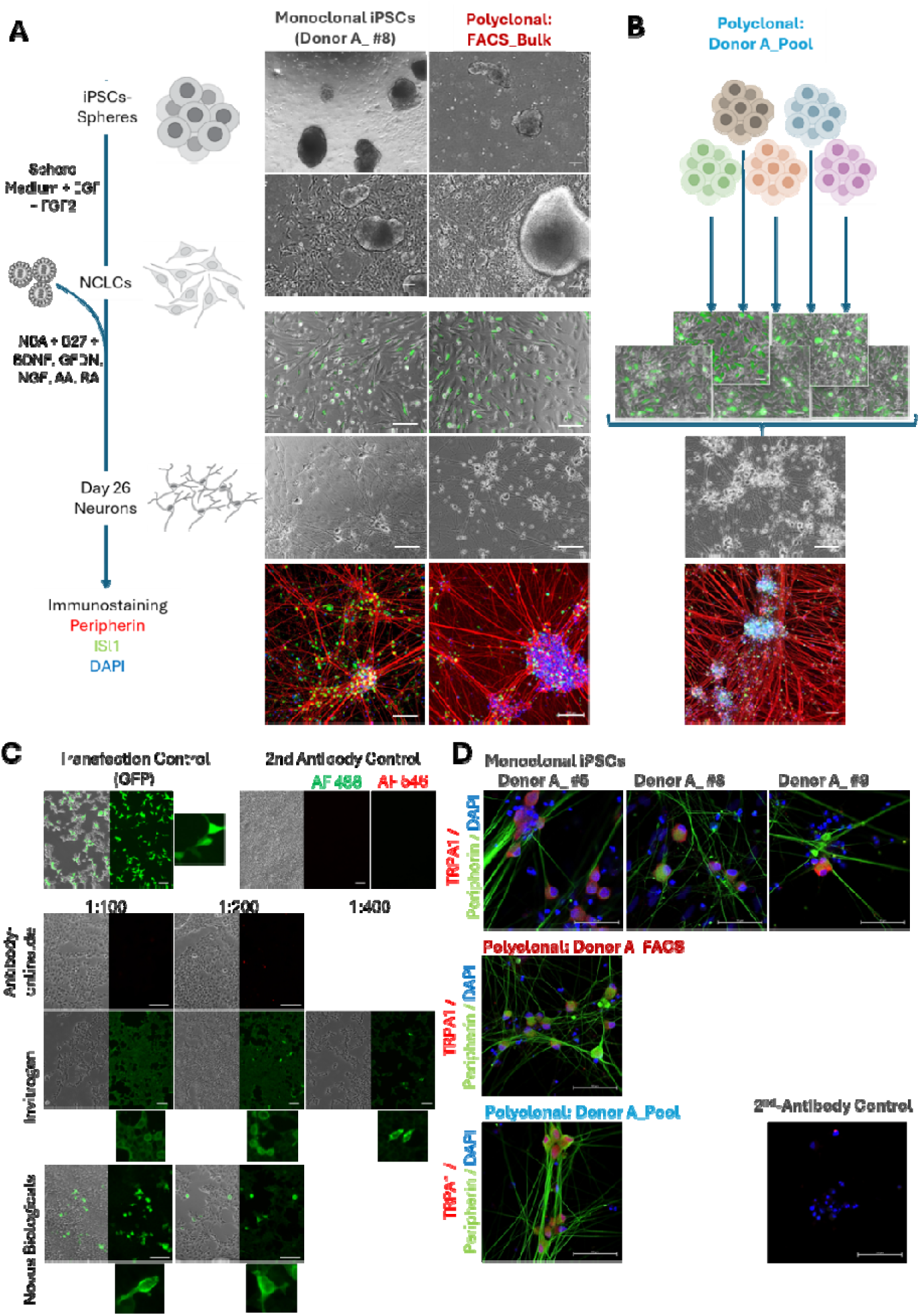
Differentiation of sensory neurons from monoclonal and polyclonal cultures. (A) Monoclonal and polyclonal iPSC lines were differentiated into sensory neurons by lentiviral-mediated ectopic expression of NEUROGENIN1 (NGN1) in neural crest-like cells (NCLCs). Left: Schematic overview of the differentiation protocol. Middle: Representative phase-contrast images of the monoclonal line Donor A-#8 and the polyclonal Donor-A_FACS-Bulk culture during differentiation. Right (B): A second polyclonal strategy was implemented by mixing five monoclonal clones per donor at day-5 of sensory neuron differentiation. All approaches yielded sensory neurons, as confirmed by immunofluorescence staining for Peripherin and ISL1. Nuclei were counterstained with DAPI. (C) Three different anti-TRPA1 antibodies were evaluated for specificity using TRPA1-transfected HEK293T cells. Transfection controls (GFP) and secondary-only controls are shown (upper panel). Each antibody was tested at two to three dilutions. For each condition, phase-contrast images, TRPA1 immunostaining, and high-magnification images are depicted. (D) Sensory neurons (FACS = d26, all others d67) derived from monoclonal and polyclonal iPSC lines were stained with Peripherin and the validated anti-TRPA1 antibody from Novis Biologicals. Nuclei were counterstained with DAPI. Scale bar: 100-µm. Schematic subimages in (A) are taken from NIH BioArt Source (Virus: NIAID Visual & Medical Arts. (10/7/2024). Virus Particle Icon. NIAID NIH BIOART Source. bioart.niaid.nih.gov/bioart/543) or from BioIcons.com (iPSCs icon by almaCELL is licensed under CC- BY 4.0 Unported https://creativecommons.org/licenses/by/4.0/, Neural_precursors icon by almaCELL is licensed under CC-BY 4.0 Unported https://creativecommons.org/licenses/by/4.0/. Neurons icon by almaCELL is licensed under CC-BY 4.0 Unported https://creativecommons.org/licenses/by/4.0/.

To generate sensory neurons, NCLCs were transduced with a lentiviral vector encoding NGN1, GFP, and a puromycin resistance cassette, together with a second vector expressing rtTA (Figure 4A). Upon doxycycline addition, NCLCs initiated GFP expression and converted into neurons expressing Peripherin and ISL1, two markers associated with sensory neuron identity. No differences in neuronal morphology or marker expression were observed between monoclonal and polyclonal cultures, indicating that polyclonal iPSC cultures are fully competent to generate sensory neurons and may serve as a suitable basis for patient-in-a-dish approaches in translational research.

Since polyclonal iPSC cultures are more susceptible to retaining parts of the Sendai virus genome, showing residual transgene expression, or containing subpopulations with aberrant genotypes or altered differentiation potential, we additionally implemented a polyclonal pool strategy during differentiation (Figure 4A). This approach enables stringent selection of high-quality monoclonal iPSC lines after reprogramming and NCLC-differentiation while generating a reduced yet representative patient-specific neuron culture. To this end, five independent iPSC lines per donor were differentiated to NCLCs and sensory neuron progenitors until day-5, the routine passaging time point. During passaging, progenitors were mixed in equal ratios to generate donor-specific pools (Donor-A_Pool, Donor-B_Pool Fig. 4B). These pooled progenitor cultures efficiently generated sensory neurons expressing Peripherin and ISL1, with no detectable differences compared to monoclonal or FACS-derived polyclonal cultures.

In summary, both monoclonal and polyclonal approaches robustly generate sensory neurons without major differences in differentiation efficiency or marker expression. These findings highlight polyclonal strategies—including bulk and pooled approaches—as promising tools for scalable, high-throughput, patient-specific disease modelling.

### 2.5 Neuron-cultures do not show TRPA1 surface expression in immunofluorescence stainings

To assess potential variability between monoclonal and polyclonal approaches, we examined the expression of the nociceptive ion channel TRPA1 by immunofluorescence staining. Since TRPA1 protein expression has not been convincingly demonstrated in human iPSC-derived neurons, we hypothesized that TRPA1 might be expressed only in a small subset of cell lines that are prone to differentiate to TRPA1-expressing neurons and TRPA1, therefore, would be more readily detectable in polyclonal iPSC-derived sensory neuron cultures.

To ensure the use of a functional antibody, we first generated a positive control by transfecting 293T cells with a TRPA1 expression plasmid. Untransfected cells present in the same dish served as negative controls for the primary antibody (Figure 4C). We tested three commercially available anti-TRPA1 antibodies (antibodies-online.de, Invitrogen, and Novus Biologicals) at two to three dilutions each. The antibodies-online.de reagent did not detect TRPA1 under any condition (Figure 4C). The Invitrogen antibody produced substantial background staining and only sporadically labeled individual cells; at higher dilutions, background was reduced. In contrast, the Novus Biologicals antibody yielded brightly fluorescent TRPA1-positive cells without generating background staining across all tested dilutions. Unlike the GFP transfection control, which labeled the entire cell, both functional antibodies primarily stained the plasma membrane and perinuclear regions, consistent with endoplasmic-reticulum–associated TRPA1 localization.

Using the validated Novus Biologicals antibody, we next examined TRPA1 expression in our iPSC-derived sensory neurons at day 26 (FACS-derived bulk and monoclonal cultures) and day 67 (monoclonal and pooled cultures) (Figure 4D, Supplementary Figure 5, 6). TRPA1 stainings presented diffusely across the whole cell area. We did not detect clear TRPA1-cell surface staining at either time point. These findings indicate that both monoclonal and polyclonal iPSC cultures are suitable for sensory neuron differentiation, but they also suggest that TRPA1 expression requires further optimization of the differentiation protocol to achieve robust surface expression.

## 3. Discussion

Monoclonal iPSC lines represent the current standard in the field, and clonal variability is typically addressed by generating and characterizing multiple independent clones per donor. However, this approach is labor-intensive, time-consuming, and costly, which poses a substantial limitation for high-throughput applications such as clinically relevant patient-specific disease modelling and drug screening. Polyclonal strategies offer a potential solution by combining multiple clones from the same donor into a single culture, thereby integrating clonal variability into all downstream experiments while markedly reducing workload, time, and cost.

Previous studies applying polyclonal reprogramming strategies have used human dermal fibroblasts as starting material and relied on episomal or Sendai virus–based reprogramming methods ^18,19^. In contrast, we implemented a polyclonal Sendai virus–mediated reprogramming strategy using peripheral blood mononuclear cells (PBMCs). PBMCs offer several advantages over fibroblasts: they are minimally invasive to obtain, commercially available reprogramming kits allow the generation of transgene-free iPSCs.

In monoclonal reprogramming, multiple clones are initially generated but only morphologically best clones survive the stringent morphological selection and undergo quality control to further select the highest quality cell lines for further application. Willmann et al. observed that polyclonal iPSC- cultures derived from dermal fibroblast do not require such enrichment steps ^18^. In contrast, we observed, that PBMC-derived polyclonal cultures also needed stringent selection methods since remaining somatic or partially reprogrammed cells overgrew iPSC-colonies. In our hands, enrichment by flow cytometry results in a polyclonal cell line that maintained pluripotency and differentiation capacity throughout all experiments.

Furthermore, FACS-based enrichment of pluripotency marker–positive cells enable the early isolation of large numbers of high-quality iPSCs. In this study, FACS and MACS enrichment were performed 34 days after transduction. This timeline could be substantially shortened in future experiments, as monoclonal colonies were isolated 20 days after transduction from the same reprogramming dish used to generate the polyclonal cultures. Those 20 days could easily be saved. Additional expansion was required only to obtain sufficient cell numbers for polyclonal enrichment at day 34. Even with this additional step, the polyclonal workflow remained faster than the monoclonal approach, which required an average of 40.8 days compared to 38 days for the polyclonal FACS-Bulk line to reach cryopreservation of ten vials for establishing a master stock. In the future, it will be advantageous to determine the earliest feasible enrichment time point that still yields clean, stable iPSC cultures with sufficient cell numbers to initiate differentiation into the somatic cell type of interest as early as possible. Such an accelerated workflow is particularly attractive for disease modelling and for subsequent personalized drug screening in commercially or clinically relevant settings, where low cost, high efficiency, scalability, and automation are essential. This approach would be highly valuable for neuropathic pain modelling were low iPSC passage numbers (passage 5–10) have been associated with improved sensory neuron differentiation efficiency ^9^, a prerequisite for efficient pain modelling and reliable drug screenings.

We assessed the quality of our iPSC cultures based on pluripotency marker expression, functional differentiation potential, genetic integrity, and Sendai virus/transgene clearance. We observed no differences between monoclonal and polyclonal cultures in pluripotency marker expression or in their ability to differentiate into all three germ layers, NCLCs, or sensory neurons. Elimination of reprogramming transgenes and Sendai virus (SeV) RNA is an important quality criterion for iPSCs, as residual transgene expression may interfere with endogenous pluripotency networks and lead to aberrant differentiation outcomes ^26–28^. In our study, all reprogramming transgenes were cleared by day 27 after initiating reprogramming. In contrast, SeV RNA persisted across several passages in monoclonal lines. Although extended culture has been reported to progressively reduce SeV levels, with viral RNA often persisting for prolonged periods ^27,29–31^. SeV expression remained stable in our cultures even after 20 passages. As expected for a polyclonal population, both FACS- and MACS-derived bulk cultures also showed consistent SeV expression. The Sendai vectors used for reprogramming lack the fusion protein (F-gene; SeV/ΔF) ^30^, rendering them non-transmissible and temperature-sensitive ^32^. Accordingly, culturing cells at 38.5 °C markedly reduced or eliminated SeV mRNA, suggesting that a short heat-treatment step could be incorporated into polyclonal reprogramming workflows to accelerate viral clearance. However, given that all reprogramming transgenes are silenced early in the process, it remains unclear whether residual SeV mRNA persists during differentiation and whether it affects lineage specification or functional maturation. Although alternative reprogramming strategies could be considered, Sendai virus–based methods remain among the most efficient and least labor-intensive approaches, making them particularly attractive for scalable or high-throughput applications ^29^.

CNV analysis revealed several aberrations that were shared across all lines from a given donor, suggesting that these variants are donor-specific rather than reprogramming-induced. Interestingly, the polyclonal iPSC line displayed fewer detectable aberrations (e.g., absence of the *KCNQ1* alteration) compared to monoclonal lines. This may reflect the fact that monoclonal iPSC lines originate from a single PBMC within a heterogeneous population. Early clonal isolation is based on colony morphology and may inadvertently enrich for cells with an advantage during reprogramming, potentially due to pre-existing or newly acquired genetic abnormalities. Such aberrant cells may be present in polyclonal cultures at very low frequencies—below the detection threshold—or may experience a proliferative disadvantage compared to other clones and undergo negative selection during polyclonal culture ^33^. Conversely, genetic aberrations can also confer a proliferative advantage, raising the possibility that polyclonal cultures may undergo clonal shifts during extended expansion. Indeed, clonal drift has been documented in iPSC cultures over time, with the number of contributing clones decreasing to fewer than ten after 38 passages in a color-barcoding system ^34^. Similarly, when three independent iPSC lines from three donors were mixed, the line with the highest doubling rate dominated the culture within only four days ^35^. Such clonal reduction is expected in polyclonal reprogramming approaches due to the heterogeneity of PBMCs and the selective pressures inherent to reprogramming and expansion. For these reasons, our polyclonal reprogramming strategy is particularly suited for experimental settings that do not require long-term expansion but instead benefit from rapid reprogramming followed by immediate differentiation.

To mitigate clonal shifts while maintaining a *patient-in-a-dish* framework, we implemented a complementary pool strategy. This approach allows the selection of high-quality monoclonal iPSC lines for each donor, followed by independent differentiation to NCLCs. Sensory neuron progenitors are then pooled in equal ratios. Since cells at this stage are not expected to proliferate, and all clones successfully differentiate into neurons, these equal ratios are likely preserved throughout differentiation. This enables the simultaneous study of multiple cell lines per patient within a single culture, while avoiding the clonal drift associated with long-term polyclonal expansion. Nevertheless, this strategy reduces workload only at the final differentiation and readout stage, not during reprogramming or early quality control.

TRPA1 plays an important role in neuropathic pain states, yet its expression has been reported only rarely in human iPSC-derived sensory neurons, and when detected, it is typically shown at the mRNA level or inferred from functional responses rather than demonstrated as protein ^21,36–38^. To address this gap, we evaluated TRPA1 protein expression using monoclonal and both polyclonal strategies (FACS-Bulk and Pool) in iPSC-derived sensory neurons. To ensure reliable detection of TRPA1, we validated three independent anti-TRPA1 antibodies using TRPA1-transfected 293T cells as a positive control. Among these, the antibody from Novus Biologicals produced the most convincing and specific staining pattern.

Despite this, we were unable to detect TRPA1 protein at the cell surface in any of the monoclonal and polyclonal iPSC-derived sensory neuron cultures. In contrast, we detected a diffuse staining of the whole cell area. Together with our polyclonal culture system—which enables the simultaneous screening of multiple clones within a single dish—these findings suggest that TRPA1 surface expression is not dependent on the iPSC-lines used but rather a limitation of the current differentiation method. If TRPA1-positive sensory neurons may arise only from specific clones with the intrinsic capacity to express TRPA1, polyclonal approaches would increase the likelihood of capturing such rare events.

Taken together, our findings support the use of polyclonal iPSC cultures for applications that require a patient-in-a-dish format without prolonged expansion, which is particularly important given the risk of clonal shifts. Their rapid generation, demonstrated compatibility with automated high-throughput workflows ^19^, and inherent integration of clonal variability enable fast, patient-specific assays. This makes polyclonal systems highly attractive for scalable drug-screening pipelines and patient-specific experimental designs.

## 4. Methods

### 4.1 Human cell lines

Peripheral blood mononuclear cells (PBMCs) were obtained from two related donors (father-daughter) suffering from a familial chronic pain syndrome characterized by widespread pain and pain hypersensitivity triggered, among others, by prolonged standing. Donor B07, female, was 11 years old at the time of blood collection; exome sequencing revealed no pathogenic or likely pathogenic variants. Donor B08, male, 36 years old at the time of blood collection, carries a TRPA1 variant (c.1743del (−)) classified as a variant of uncertain significance by the human genetics department, Uniklinik RWTH Aachen, Germany. All procedures were conducted following informed consent as approved by the local ethics committee at Uniklinik RWTH Aachen (EK 243 – 18).

### 4.2 PBMC isolation

PBMCs were isolated by density-gradient centrifugation: Whole blood was passed through a 70 µm strainer, diluted 1:1 with D-PBS (Gibco) containing 2 mM EDTA (Gibco), and carefully layered onto PanColl (PanBiotech 1,077g/ml). After centrifugation at 751 × g for 20 min without brake, the PBMC layer was collected, washed twice with Ca²⁺/Mg²⁺-free D-PBS + 2 mM EDTA, and pelleted at 423 × g and 300 × g, respectively. Cells were resuspended in Ca²⁺/Mg²⁺-free D-PBS + 2 mM EDTA, counted using Trypan Blue exclusion, and subjected to an additional low-speed centrifugation step. For cryopreservation, PBMCs were resuspended in 90% FBS and 10% DMSO (both Sigma Aldrich), aliquoted at approximately 5 × 10 cells per vial, cooled at −80 °C in a controlled-rate freezing container overnight, and transferred to the vapor phase of liquid nitrogen for long-term storage.

### 4.3 Reprogramming of PBMCs to iPSCs

Reprogramming was performed using the CytoTune™-iPS 2.0 Sendai virus Kit (Thermo Fisher Scientific) following the manufacturers recommendations for feeder-free reprogramming of PBMCs with slight modifications. Prior to transduction, PBMCs were thawed and cultured for 4 days in PBMC Medium (StemPro™-34 medium (Gibco) supplemented with human recombinant 100ng/ml SCF (Life Technologies), 100ng/ml FLT3L (Life Technologies), 20ng/ml IL-3 (Life Technologies), and 20ng/ml IL-6 (Peprotech) with an initial seeding density of 5×10^5^ cells /ml with one ml / 24 well of an ultra-low attachment plate. 50% of the medium was replaced daily. To initiate the reprogramming, cells were collected and 5×10^5^ cells were mixed with all 3 Sendai viruses with and MOIs of KOS = 5, Myc =5, hKlf4=3 in a final volume of 1ml of PBMC-medium. Cell/Virus mix was centrifuged for 30min at 1000g and subsequently seeded to one well of a 24-wellplate with an additional ml of PBMC-medium. The next day, cells were centrifuged to remove virus, resusupendend in fresh PBMC- medium and seeded back to the same, washed 24 well. On day 4 post-transduction, cells were transferred to Vitronectin-coated 10cm-plates in PBMC-Medium without cytokines at a density of 1×10^4^-1×10^5^ cells/plate. Emerging colonies were monitored daily, and 50% medium was exchanged every 24 h. At d7, 50% spend medium was replaced with iPS brew medium (Miltenyi Biotech) followed by complete medium changes daily until isolation of presumptive iPSCs.

### 4.4 Clonal selection of iPSCs

Individual colonies with typical pluripotent morphology—compact appearance, high nucleus-to-cytoplasm ratio, and sharp colony borders—were manually isolated between days 18–25 post-transduction. iPSC-colonies were identified with an Evos M7000 Imaging System (Life Technologies) placed under the cell culture bench. Cells were scratched off the surface and collected with a 200µl pipette tip and transferred into individual Vitronectin-coated 24-wells containing iPS- brew medium supplemented with 10 µM ROCK inhibitor (Y-27632) for the first 24 h. Clonal lines were expanded for cryopreservation and further experiments.

### 4.5 Isolation of iPSCs for polyclonal cultures

For polyclonal reprogramming, all remaining cells after clonal selection within a dish were maintained and passaged ones. To enrich for reprogrammed cells, presumptive iPSC were (1.) routinely passaged or (2.) purified by flow cytometry (FACS) or (3.) magnetic activated cell sorting (MACS). For MACS and FACS, cells were dissociated into single cells using Accutase (Sigma Aldrich) for 5min at 37°C and counted with a CellDrop (DeNovix). 50% of the cell suspension was stained with fluorophore-conjugated anti-TRA-1-60-PE and anti-SSEA4-APC antibodies (Supplementary Table 1). Staining was performed in iPS-brew medium for 30min at 4°C. All cell fractions were sorted on Becton Dickinson (BD) FACS Aria Fusion and replated onto Vitronectin coated 6-Wells in iPS-brew Medium supplemented with 10µM ROCK (Stem Cell Technologies). MACS purification followed the manufacturer’s manual provided with Anti-TRA-1-60 MicroBeads (Miltenyi Biotech) Cells were passed through MS columns. The positive and negative fraction was eluted, plated onto Vitronectin-coated 6-Wells in iPS brew supplemented with 10µM ROCK, and expanded under standard iPS-culture conditions.

### 4.6 iPS-Culture and Cryopreservation

iPSCs were maintained under feeder-free conditions on recombinant human vitronectin (VTN-N, Life Technologies) and cultured in iPS Brew medium (Miltenyi Biotec). Coating and medium preparation followed the manufacturer’s instructions. Cultures were supplied with fresh medium daily with no medium changes the day after passaging and weekends. Once colonies reached approximately 70% confluence or displayed dense central regions, cells were passaged using 0.5 mM EDTA in PBS at a split ratio of 1:6 to 1:10. For cryopreservation, detached cells were resuspended in freezing medium consisting of 90% iPS brew medium, 10% DMSO, and 10 µM Y-27632, and transferred to a controlled rate freezing container (Mr. Frosty™) for overnight cooling at −80 °C. Vials were subsequently stored in the vapor phase of liquid nitrogen. All iPSC lines were routinely screened for mycoplasma contamination using the MycSPY kit (Biontex).

### 4.7 Immunofluorescence Staining

For immunofluorescence stainings, cells were seeded on glas coverslips and routinely cultured until experiment. Cells were fixed in 4% paraformaldehyde (Sigma Aldrich) for 15 min at room temperature, permeabilized with 0.1% Triton X-100 (Sigma Aldrich), and incubated in 5% donkey serum (Pan Biotech) to block nonspecific binding. Primary antibodies (Supplementary Table 2) were applied overnight at 4 °C. After three washes with PBS, cells were incubated with species-appropriate secondary antibodies (Supplementary Table 3) and counterstained with DAPI (Gibco; NucBlue Fixed Cell ReadyProbes Reagent). Fluorescence images were acquired using a LSM M700 confocal microscope (Zeiss).

### 4.8 Fluorescent Activated Cell Sorting

For flow cytometry, iPSCs were dissociated into single-cell suspensions using Accutase (PanBiotech), pelleted by centrifugation, and resuspended in FACS buffer (5% FBS, 95% PBS). Cell numbers and viability were determined using a CellDrop automated counter (DeNovix) with Trypan Blue exclusion. A total of 1×10 cells were incubated with primary antibodies diluted in 100 µL FACS buffer for 30– 60 min at 4 °C (antibodies are listed in supplementary table 1). After two washes with PBS, cells were resuspended in FACS buffer for acquisition. Measurements were performed on a BD FACS Canto II, and data were analyzed using FCS Express 7 (DeNovo Software). Cells were first gated based on forward and side scatter properties, followed by definition of positive and negative populations using isotype controls with a background fluorescence threshold of 0.5%. Graphs were exported directly from the analysis software.

### 4.9 Sendai virus and transgene clearance

Total RNA was isolated using the NucleoSpin RNA Mini Kit (Macherey-Nagel) following the manufacturer’s protocol. RNA concentrations were determined with a NanoDrop 2000c spectrophotometer, and samples were stored at −80 °C until further use. Reverse transcription was performed using the SensiFast cDNA Synthesis Kit (Meridian Biosciences) according to the manufacturer’s instructions. For each reaction, 1 µg RNA was used as input. cDNA was stored at 4 °C for short-term use or at −20 °C for long-term storage. To assess clearance of residual Sendai virus and exogenous reprogramming factors, PCR was carried out using primers targeting Sendai viral RNA and the transgenes encoding KLF4, OCT3/4, SOX2, and c-MYC (all Primer Sequences are listed in supplementary table 4). Reactions were prepared with OneTaq PCR Master Mix (New England Biolabs) containing 0.1 µM of each primer, 0.2 mM dNTPs, and either 125 ng cDNA (for SeV, KLF4, c-MYC) or 250 ng cDNA (for KOS). Amplification was performed on a thermal cycler (Peqstar; VWR) using the following program: 94 °C for 5 min; 35 cycles of 94 °C for 30 s, 55 °C for 45 s, and 68 °C for 40 s; followed by a final extension at 68 °C for 5 min. Positive and negative controls were included in every run. PCR products were analyzed by agarose gel electrophoresis. A 2% agarose gel was prepared in 1× TAE buffer and supplemented with RedSafe for DNA visualization. Ten microliters of each PCR reaction and 5 µL of a 100 bp DNA ladder were loaded directly onto the gel. Electrophoresis was performed at 90 V for 28 min, and DNA bands were visualized under UV illumination using the Chemostar Touch imaging system (Intas Science Imaging Instruments). Band intensities were quantified in Fiji. For each lane, a rectangular ROI was placed around the PCR band of interest, and a second ROI was positioned directly above or below the band within the same lane to capture the local background. Integrated density values (Mean × Area=IntDen) were obtained for both ROIs, and background-corrected signal intensities were calculated as InDen_corr_=IntDen_GOI_−(Mean_background_ x Area_GOI_). Background corrected values were normalized to the corresponding background-corrected HPRT signal from the same lane to account for differences in DNA input and PCR efficiency.

### 4.10 Copy number variation (CNV) analysis

iPSC pellets were submitted to Life & Brain GmbH (Bonn, Germany) for genomic DNA isolation and microarray-based quality control. The “Package 2 – Basic Report” workflow was performed, which includes CNV partition analysis using an Illumina microarray platform comprising approximately 700,000 genotyped markers. Results file was provided by Life&Brain, Bonn (Germany). For downstream evaluation, sex chromosomes were omitted from CNV interpretation due to methodological limitations in accurately determining copy number states on X and Y chromosomes (Panopoulos et al., 2017). CNV-List included genomic coordinates (chromosome, start, end), estimated copy number values (0–4), and associated confidence scores. CNV segments were classified based on their copy number state, where values of 0 and 1 indicate deletions, 2 represents diploid copy number, and values ≥3 indicate copy number gains or amplifications. To focus on biologically relevant events, segments corresponding to the diploid range (copy number = 2; defined as “CNV Bin: Min 1.5 to Max 2.5”) were excluded from downstream CNV analyses, except in cases annotated as loss of heterozygosity (LOH). LOH regions were identified based on annotation in the “Comment” field and retained as a separate category, as these represent copy-neutral allelic imbalance events. Genomic coordinates were standardized and processed using custom Python scripts (Python 3.x, pandas, NumPy, matplotlib). Chromosomal positions were converted into cumulative genomic coordinates to enable genome-wide visualization. CNV sizes were calculated from segment boundaries, and values were scaled for graphical representation. Confidence scores extracted from annotation fields were used to modulate point transparency in visualization. Gene annotation of CNV regions was performed using the Ensembl human gene annotation (GRCh38). Gene transfer format (GTF) files were parsed to extract gene coordinates, and overlap between CNV segments and gene loci was determined using interval-based matching implemented in Python. A gene was considered affected if any overlap with a CNV region was detected. For functional interpretation, CNV-associated genes were compared against a curated gene list derived from Gene Ontology (GO) terms relevant to pluripotency, neuronal differentiation, neural crest identity, glial biology, and pain-related pathways. The list was extended by selected genes published by Ghosh & Som, 2020; Tavares-Ferreira et al., 2022; Körner et al., 2024.

### 4.11 Trilineage differentiation

hiPSCs were cultured to approximately 70% confluency. For initiation of three-germ-layer differentiation, cells were washed with PBS, incubated with 1 mL 0.5 mM EDTA in PBS for 3 min, centrifuged (100 × g, 30 s) and resuspended in 2 mL medium composed of 50% DMEM/F-12 and 50% iPS-Brew. Cells were transferred to ultra low attachment plates in the presence of 10 µM Y-27632. After 24 h, medium was carefully exchanged by 2 mL trilineage differentiation medium (80% DMEM-F12 + Glutamax, 20% FBS (Sigma Aldrich), 1x MEM-NEAA, 0,2% 2-Mercaptoethanol (all Gibco)). Medium was exchanged every 2–3 days. On day 7, Embroyd Bodies were collected and plated onto Geltrex-coated 6-well plates. Medium was replaced every 2–3 days. After 14 days of differentiation, cells were detached using 0.5 mL Accutase, pelleted, and stored at −80 °C until further processing. RNA was isolated from cell pellets collected before and after trilineage differentiation and reverse-transcribed into cDNA as described in the section Sendai virus detection. For each time point, a no-reverse-transcriptase (no-RT) control was included. Each qPCR run was performed in triplicates. All primer sequences are listed in supplementary table 4. cDNA samples were diluted to a final concentration of 2 ng/µL in nuclease-free water. Primer-specific master mixes were prepared containing 0,25µ of each primer, 6 µL SensiMix SYBR No-ROX (Bioline, QT650-05), and 0.35 µL nuclease-free water per reaction. For each well, 7 µL of master mix and 5 µL of template (cDNA, no-RT, or no-template control) were dispensed into 96-well plates and sealed. qPCR was performed on a Bio-Rad CFX system using the following cycling protocol: 95 °C for 10 min; 40 cycles of 95 °C for 15 s and 60 °C for 1 min; followed by a final denaturation at 95 °C for 1 min. Melt-curve analysis was conducted from 60 °C to 95 °C in 0.5 °C increments to verify amplification specificity. Relative gene expression was calculated using the ΔΔCt method, normalizing target gene expression to GAPDH and HPRT and comparing differentiated samples to undifferentiated controls. Data extraction was performed using BioRad CFX Maestro 2.2 Software (Bio-Rad, Version 5.2.008.0222) and plotted in python.

### 4.12 Differentiation of neural crest-like cells and sensory neurons

Sensory neuron differentiation was performed following Schrenk-Siemens et al. 2022 with minor adaptations ^21^. iPSCs were expanded in 10 cm dishes and detached at 70–80% confluency using the routine EDTA-based passaging procedure. Cell clusters were transferred to uncoated 10 cm dishes (Greiner) containing 10 mL sphere medium (50% DMEM/F12+GlutaMax, 50% Neurobasal, 0,5% GlutaMax, 0,5x B27-Supplement, 0,5x N2-Supplement, 1x Penicillin/Streptomycin (all purchased from Gibco, 10ng/ml hEGF, 10ng/ml hFGFbasic (Peprotech)) supplemented with 10 µM Y-27632 (StemCell Technologies). From day 1 onward, medium was replaced every 2–3 days without Y-27632. Spheroids typically attached within 6–10 days, after which neural crest like cells (NCLCs) began to migrate outward. Once a dense ring of NCLCs had formed, floating spheroids were collected and transferred to new dishes to generate additional outgrowth. Attached spheroids were removed by aspiration, and remaining NCLCs were detached with Accutase (Sigma Aldrich), centrifuged (200 × g, 5 min), and counted using a CellDrop (DeNovix) counter with Trypan Blue. NCLCs were cryopreserved in sphere medium containing 10% DMSO and 10 µM Y-27632.

For further differentiation, NCLCs were thawed two days before transduction. A total of 1 × 10 cells/cm² was plated on Geltrex-coated (Gibco) dishes in sphere medium. If confluency remained below 50%, additional NCLCs were thawed the following day to ensure sufficient density. Cells were transduced with two lentiviral vectors: one encoding NGN1, GFP, and a puromycin resistance cassette under Tet-On control, and a second encoding rtTA. Sphere medium was supplemented with 10 mM HEPES and 8 µg/mL protamine sulfate (Sigma Aldrich) for transduction, which was performed at an MOI of 0.75. After 24 h, cells were washed three times with PBS and cultured in sphere medium containing 2 µg/mL doxycycline (Sigma Aldrich) marking day 0 of sensory neuron differentiation. From day 1 to day 10, all media contained 2g/ml Dox and were changed daily. At d1, cells transitioned to sensory neuron differentiation medium (50% DMEM/F12+GlutaMax, 50% Neurobasal, 0,5% GlutaMax, 0,5x B27-Supplement, 0,5x N2-Supplement, 1x Penicillin/Streptomycin (all purchased from Gibco, 20ng/ml hBDNF, 10ng/ml hGDNF, 10ng/ml hNGF (Peprotech)) and on day 3, 10 µg/mL puromycin was added for 24 h to enrich for transduced cells. On day 5, cells were detached with Accutase, counted, and seeded onto coverslips coated sequentially with 15 µg/mL poly-L-ornithine and 10 µg/mL laminin/fibronectin. All coating steps were performed overnight at 37 °C. From day 5 onward, 50% media chaned were performed. Starting from d8, cells were gradually transitioned to sensory neuron maturation medium (100% Neurobasal A without Glucose/NaPyruvate, 1x B27-Supplement, 1x N2-Supplement, 1x GlutaMAX, 5 mM Glucose, 0.055 mM NaPyruvate, 0.125 µM Retinoic Acid, 0.2 mM Ascorbic Acid, 0.5 µg/mL Laminin (all Sigma Aldrich), 35 mM NaCl, 20 ng/mL BDNF, 20 ng/mL GDNF, 20 ng/mL NGF (Peprotech)). Beginning on day 11, medium was changed every other day without doxycycline. At this stage, non-neuronal proliferation was suppressed by adding 2 µM cytosine arabinoside (Ara-C) for 16–20 h, followed by complete medium exchange. Medium changes continued every other day until the cells were used for experiments.

### 4.13 Virus Production

HEK293T/17 cells (ATCC CRL-11268) were maintained in high-glucose DMEM supplemented with 10% FBS (Sigma-Aldrich) and 1× sodium pyruvate (Gibco) (293t-Medium). Cells were passaged two to three times per week using 0.05% Trypsin/EDTA (Gibco) at split ratios of 1:10 to 1:20. For virus production, 3×10^6^cells were seeded per 10cm dish in 293t-Medium one day prior to transfection. Lentiviral vectors for co-expression of NGN1, eGFP, and a puromycin resistance cassette under Tet-On control, or with rtTA, were generated by calcium phosphate–mediated co-transfection of the helper plasmids pRSV-REV (Addgene #12253), pMD2.G (Addgene #12259), and pMDLg/pRRE

(Addgene #12251) with either FUW-rtTA (Addgene #20342) or FUW-TetO-Ngn1-P2A-EGFP-T2A-Puro at a ratio of 1:1:2:3. All plasmids were obtained from Katrin Schrenk Siemens (Institute of Pharmacology, University of Heidelberg). Sixteen to twenty hours after transfection, the medium was replaced with virus production medium (293t-Medium supplemented with 10 mM HEPES). Viral supernatants were collected at 48 h and 72 h post-transfection. Supernatants containing NGN1 and rtTA lentiviruses were mixed and precipitated by addition of 80 µg/mL polybrene and 80 µg/mL chondroitin sulfate (Sigma-Aldrich), followed by centrifugation at 4000 rpm for 20 min. Pellets were resuspended in 1% of the initial volume using DMEM supplemented with 20 mM HEPES. Concentrated viral stocks were stored at −80 °C. Viral titers were determined by transducing 1 × 10⁵ HEK293T cells with defined volumes of virus. Twenty-four hours after induction of eGFP expression with 2 µg/mL doxycycline (Sigma-Aldrich)the percentage of eGFP-positive cells was quantified by flow cytometry (BD FACSCanto).

### 4.13 Transfection of 293t cells with TRPA1-encoding plasmid

HEK293T cells were maintained in high-glucose DMEM supplemented with 10% FBS 37 °C and 5% CO. For transfection, cells were seeded one day prior at 5×10 cells/cm² and reached approximately 50% confluency at the time of transfection. Cells were transfected with 1.5 µg total plasmid DNA pCDNA3.1/V5-His-hTRPA1 ^39^. DNA was diluted in 50 µL NaCl. 1,5µl jetPEI (Peqlab) was diluted in 48,5µl NaCl, and 50 µL jetPEI was mixed with 50 µL DNA solution to form DNA– jetPEI complexes. Complexes were incubated for 20 min at room temperature and added dropwise to the cells. One day later, cells were fixed and immunostained following the immunocytochemistry protocol described above to evaluate the performance of three commercial TRPA1 antibodies.

## Author contributions

AN conceived and supervised the study, interpreted the results and wrote the manuscript.

LR performed experiments, analysed the data and interpreted the results.

PH carried out the HEK293T transfections for TRPA1 antibody testing and assisted in establishing the PCR assays for detection of residual Sendai virus and reprogramming transgenes.

MFD and NWMvdB contributed to the acquisition and interpretation of data.

AS and MH recruited the patients and performed blood sampling.

AL and KE performed genetic evaluation of the donors.

ALa conceived and supervised the study, interpreted the results, provided funding and contributed to manuscript writing.

All authors reviewed and approved the final manuscript

## Acknowledgements

This work was supported by the Flow Cytometry Facility of the Interdisciplinary Center for Clinical Research (IZKF) within the Faculty of Medicine at RWTH Aachen University.

## Funding

ALa, MFD, and MH were supported by a grant from the Interdisciplinary Center for Clinical Research within the Faculty of Medicine at the RWTH Aachen University (IZKF TN1-1/IA 532001, IZKF TN1-9/IA 532009). ALa has received funding from the DFG, German Research Foundation 363055819/GRK2415, DFG, German Research Foundation 368482240/GRK2416, DFG, German Research Foundation LA 2740/6-1, via the BMBF consortium “Bio2Treat” (German Federal Ministry of Education and Research / Bundesministerium für Bildung und Forschung, BMBF, “Chronische Schmerzen- Innovative medizintechnische Lösungen zur Verbesserung von Prävention, Diagnostik und Therapie”, contract number 13GW0334B, and via the BMBF consortium “Precision2Treat” (German Federal Ministry of Education and Research / Bundesministerium für Bildung und Forschung, BMBF, “Patientenindividuelle Therapie von Schmerzpatienten mittels induzierter pluripotenter Stammzellen (Precision2Treat)”, contract number 13GW0656.

## Data availability statement

All data are available upon request.

## Competing Interests

MFD reports grants from Pfizer Pharmaceuticals (ASPIRE 2018); consulting fees from Pfizer, Alnylam, Akcea, AstraZeneca, Amicus Therapeutics, Applied Therapeutics, and Sobi; payment or honoraria from Pfizer, Alnylam, Akcea, Amicus Therapeutics, AstraZeneca, and Sobi; support for attending meetings and/or travel from Pfizer, Alnylam, Akcea, AstraZeneca, Sobi, and Amicus Therapeutics; participation on advisory boards for Pfizer, Alnylam, Akcea, AstraZeneca, and Purpose Pharma; and a leadership role in the European CMT Research Association (ERCA).

ALa receives councelling fees from Grünenthal, Netri and Orion.

## Additional Information

AI assistance (ChatGPT, accessed august 2026) was used to generate initial code templates for data processing and analysis. Final code reflects substantial manual revision. We used Microsoft Copilot (version accessed in March 2026) to support text revision. All outputs were reviewed, verified, and adapted by the authors.

