## Supplementary data for "Polyclonal sensory neuron derivation from iPSCs as an efficient alternative to single clone strategies for pain-relevant *in vitro* models"

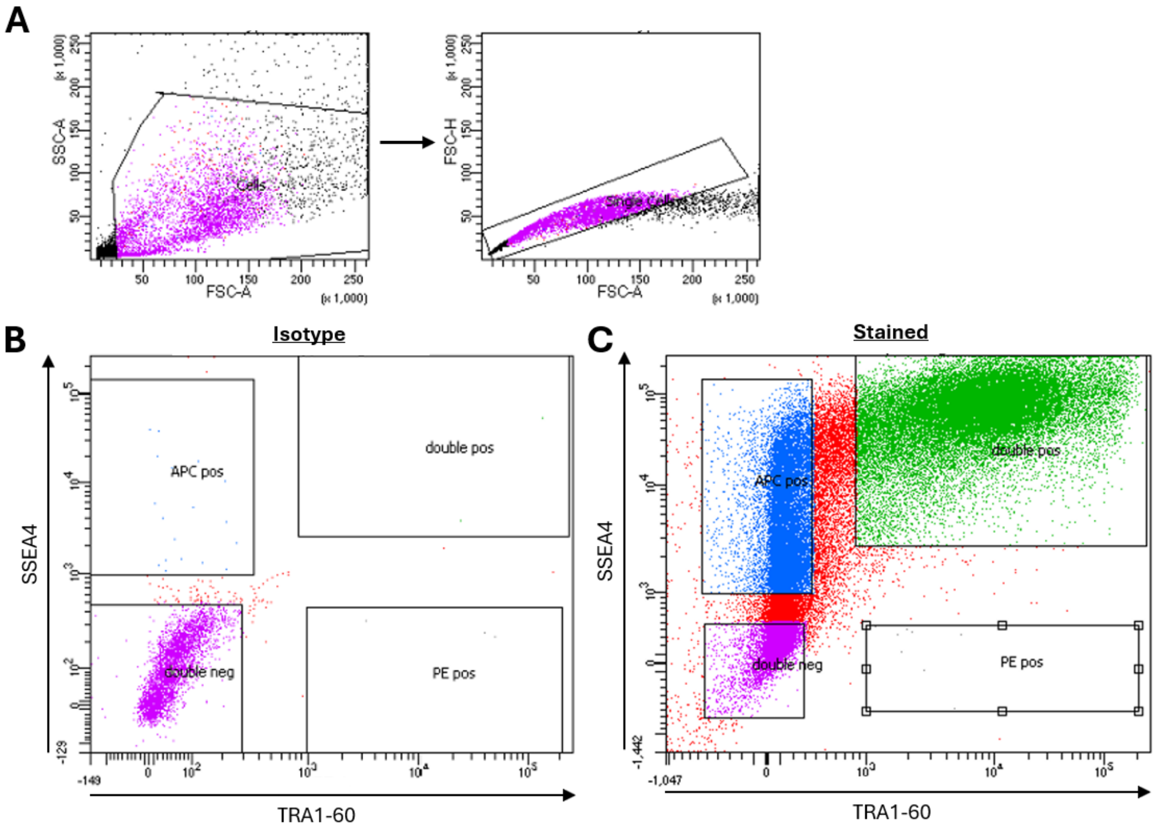


**Supplementary Figure 1 FACS sorting of polyclonal cultures of Donor_A.**

(A) Cells were first gated based on forward and side scatter (FSC/SSC) properties, followed by gating on single cells (FSC-A/FSC-H). (B) Sorting gates were defined using isotype control stainings to establish background fluorescence. (C) Marker-positive populations were subsequently sorted according to TRA-1-60 and SSEA4 expression.

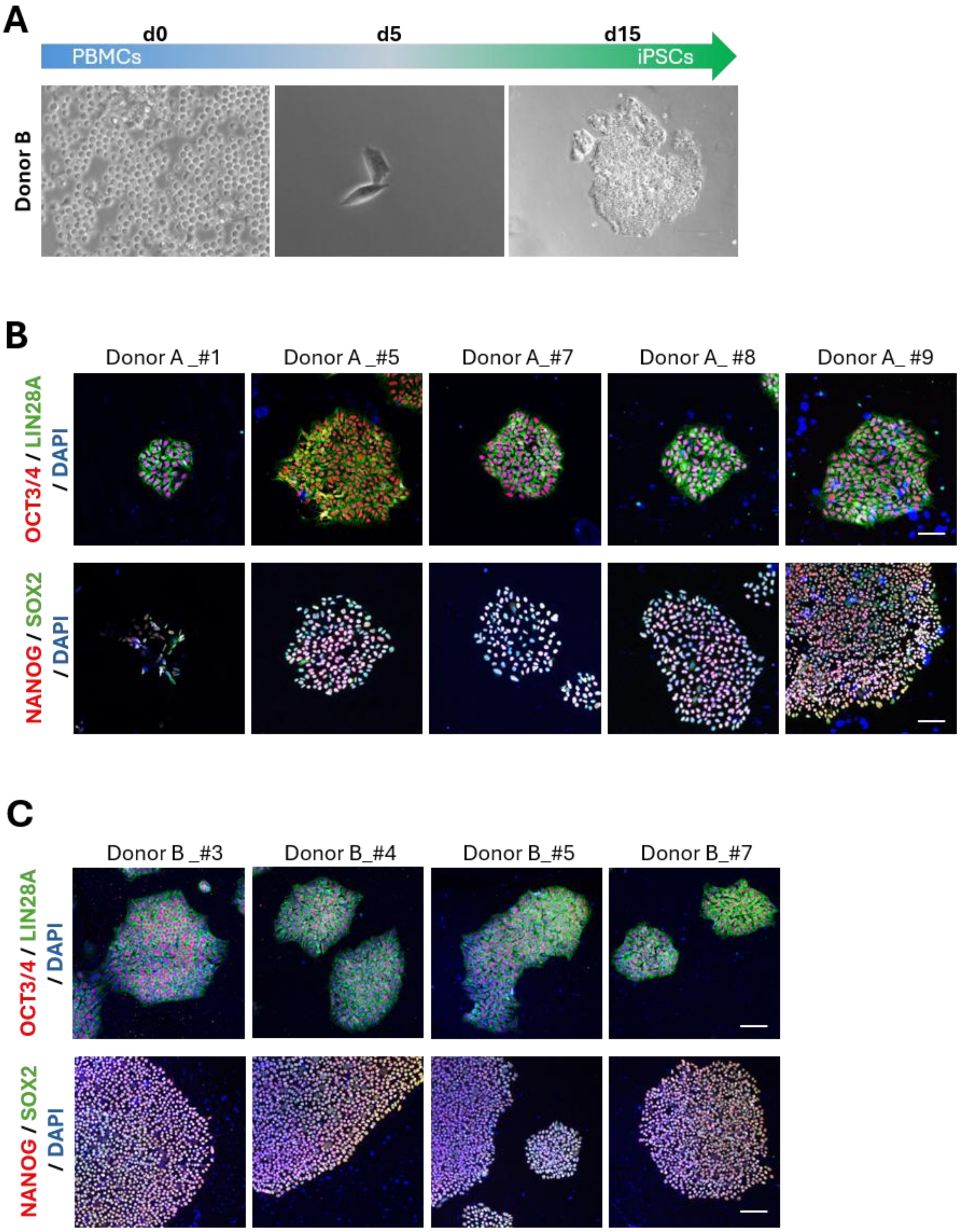


**Supplementary Figure 2 Reprogramming of Donor B and detection of pluripotency-associated markers by immunofluorescence.**

(A) Following Sendai virus transduction (d0), PBMCs from Donor B progressively acquired iPSC-like morphology over the course of 15 days, comparable to the reprogramming dynamics observed in Donor A. (B, C) Immunofluorescence staining confirmed robust expression of the core pluripotency markers OCT4, SOX2, NANOG and LIN28A in all monoclonal iPSC lines derived from Donor B as well as in representative lines from Donor A. Scale bar: 100 μm.

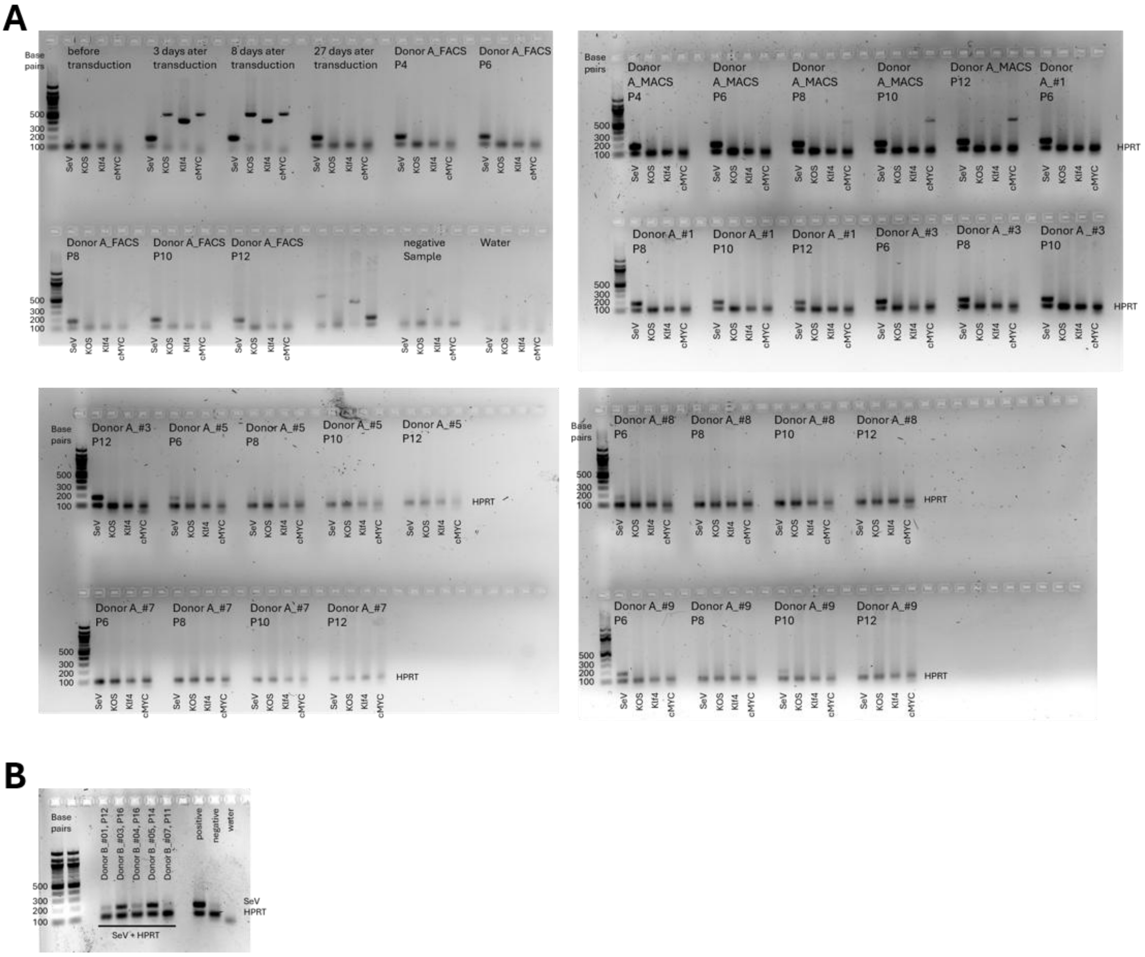


**Supplementary Figure 3 Agarose Gels to detect Sendai-Virus (SeV) and reprogramming transgene expression.**

(A) Complete, inverted agarose gel images corresponding to the PCR analyses presented in Figure 3 are shown. All amplicons for Sendai virus (SeV, 181bp) and the reprogramming transgenes KOS (528bp), KLF4 (410bp) , cMYC(532bp) are shown alongside the HPRT loading control (102 bp). Lanes include samples collected during the reprogramming time course as well as monoclonal and polyclonal iPSC lines derived from Donor A. (B) Detection of SeV in Donor B in established monoclonal iPSC lines.

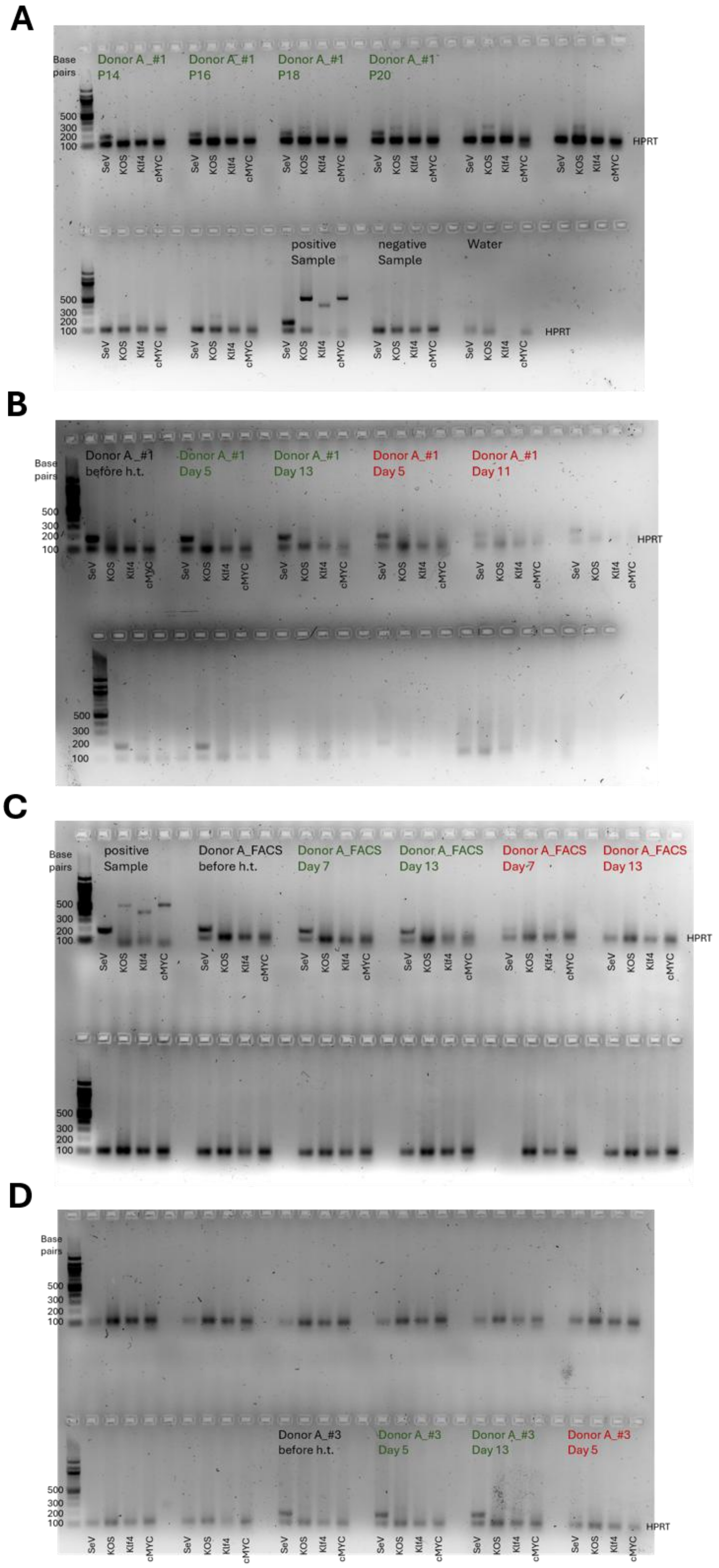


**Supplementary Figure 4 Complete, inverted agarose gel images after prolonged culture and heat treatment.**

Complete, inverted agarose gel images corresponding to the PCR analyses presented in Figure 3 G-I are shown. All amplicons for Sendai virus (SeV, 181bp) and the reprogramming transgenes KOS (528bp), KLF4 (410bp) , cMYC(532bp) are shown alongside the HPRT loading control (102 bp). Lanes include samples collected during prolonged culture (A) as well as monoclonal and polyclonal iPSC lines derived from Donor A cultures at 38,5°C for 5 days (red) or 37°C (green).

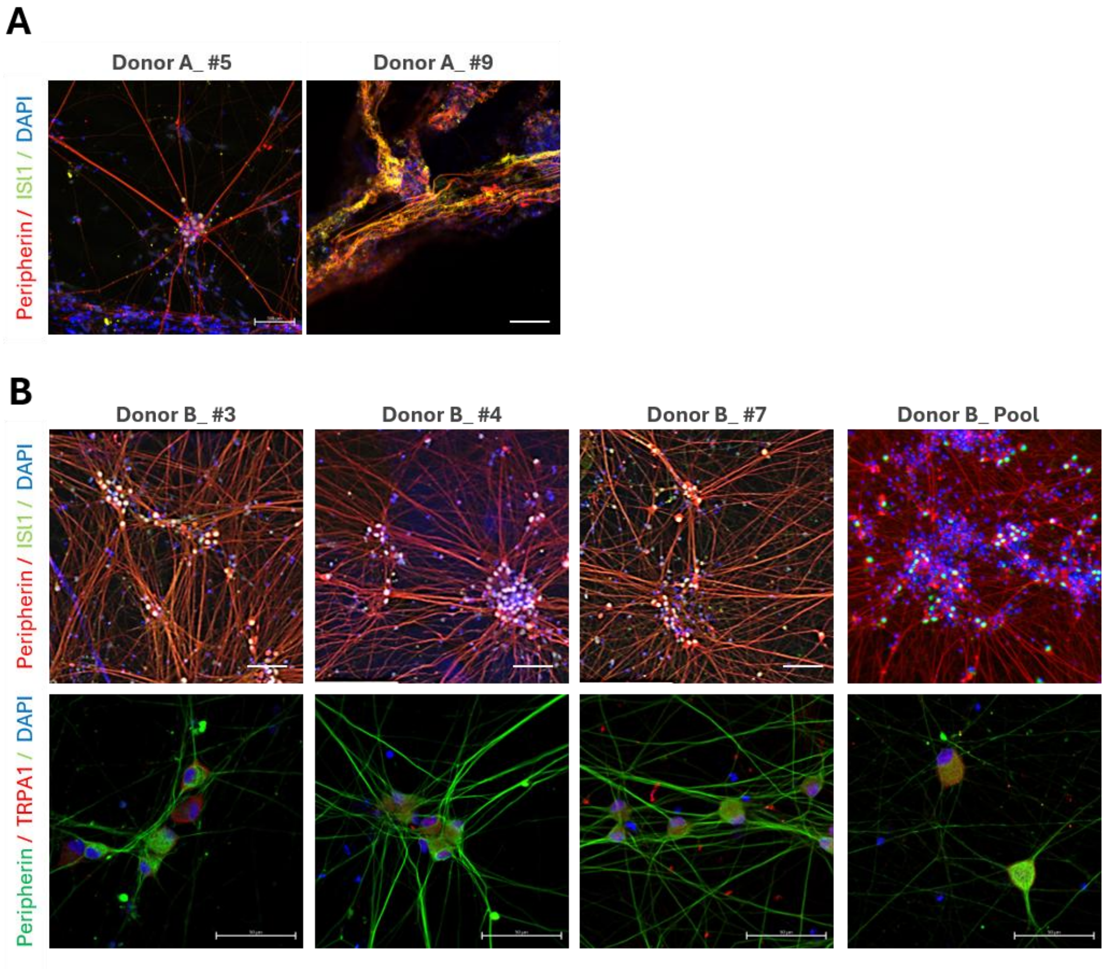


**Supplementary Figure 5 Immunofluorescence Staining of monoclonal and polyclonal iPSC-derived neurons.**

Immunofluorescence analysis of all remaining monoclonal and polyclonal iPSC-derived sensory neuron cultures of Donor A (A) and Donor B (B) not shown in Figure 4. Cells were stained for Peripherin and ISL1 or Peripherin and TRPA1 and nuclei were counterstained with DAPI.

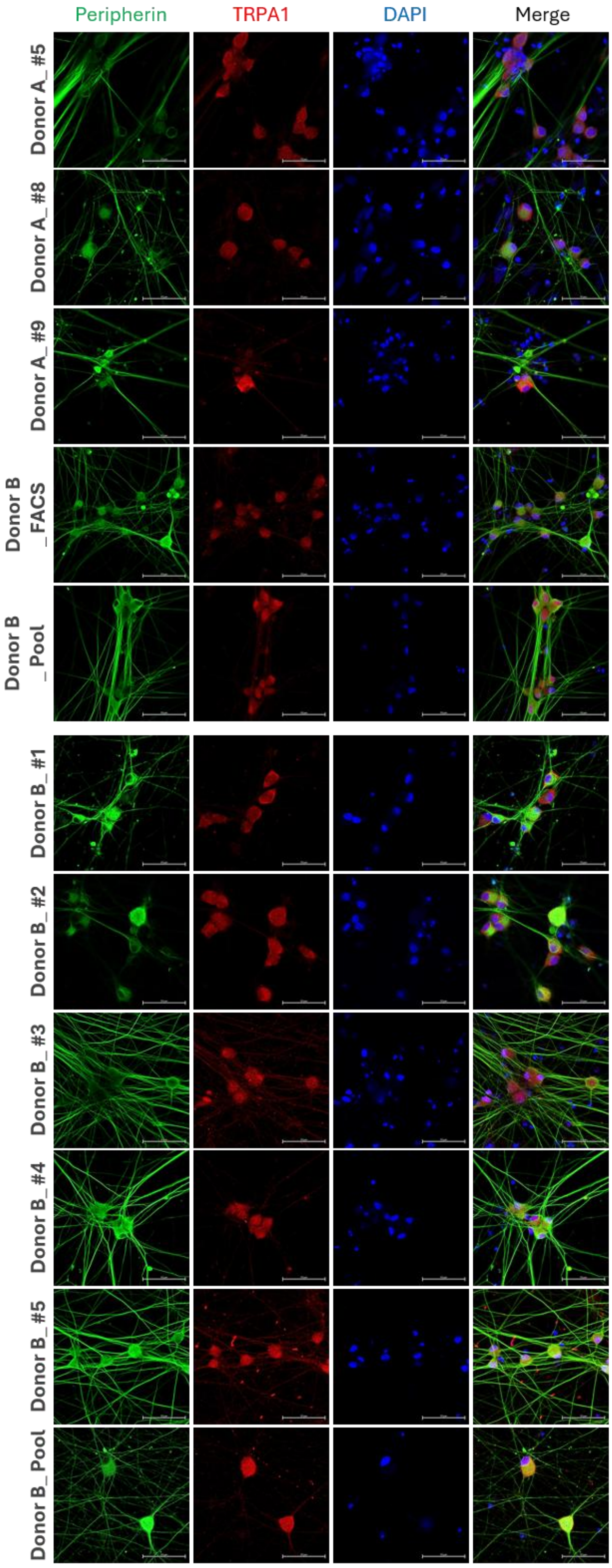


**Supplementary Figure 6 Single-channel images of Peripherin- and TRPA1-stained sensory neurons.**

Neurons derived from monoclonal and polyclonal iPSC cultures were stained for Peripherin (green), TRPA1 (red), and nuclei (DAPI, blue). Single-channel and merged images are shown for each donor culture. TRPA1 staining appeared diffusely distributed throughout the cytoplasm without clear membrane localization in all cultures. Scale bar = 50 μm.

**Supplementary Table 1 Antibodies for fluorescent activated cell sorting and magnetic activated cell sorting**

| antibody | fluorochrome | manufacturer | Ordering Number | dilution |
|---|---|---|---|---|
| anti-human TRA1-60-R | PE | Bio Legend | 330610 | 1:500 |
| PE mouse IgM, isotype control | PE | Bio Legend | 401609 | 1:500 |
| anti-human SSEA4 | Alexa Fluro 647 | Bio Legend | 330408 | 1:500 |
| mouse IgG3, isotype control | Alexa Fluro 647 | Bio Legend | 401321 | 1:500 |
| Anti-TRA-1-60 MicroBeads, human | / | Miltenyi Biotec | 130-100-832 | 20ul/2x$10^6$ cells in 100µl |

**Supplementary Table 2 Primary Antibodies**

* exact usage of the antibody needs to be evaluated further

| primary antibody | host | manufacturer | Ordering Number | dilution |
|---|---|---|---|---|
| Sox2 | mouse | Cell Signaling Technologies | 4900 | 1:400 |
| Nanog | rabbit | Cell Signaling Technologies | 4903 | 1:200 |
| Lin28A | mouse | Cell Signaling Technologies | 5930 | 1:800 |
| Oct3/4 | rabbit | Santa Cruz | SC-9081 | 1:500 |
| ISL1 | rabbit | Synaptic Systems | 406003 | 1:2000 |
| Anti-beta III Tubulin | chicken | Novus Biological | NB100-1612 | 1:1000 |
| Peripherin | mouse | Santa Cruz | SC-377093 | 1:500 |
| TRPA1 | rabbit | Novus Biological | H00008989-D01P | 1:200 |
| TRPA1 | sheep | antibodies-online.de | ABIN378598 | * |
| TRPA1 | rabbit | Invitrogen | PA1-46159 | 1:400 |

**Supplementary Table 3 secondary antibodies**

| secondary antibody | manufacturer | Ordering Number | dilution |
|---|---|---|---|
| Alexa fluor™ 555 goat anti-rabbit IgG (H+L) | Invitrogen | A-21429 | 1:1000 |
| Alexa fluor™ 488 goat anti-mouse IgG (H+L) | Invitrogen | A-11001 | 1:1000 |
| Alexa fluor™ 633 goat anti-mouse IgG (H+L) | Invitrogen | A21126 | 1:1000 |
| Alexa fluor™ 488 goat anti-chicken IgG (H+L) | Invitrogen | A11039 | 1:1000 |
| Alexa fluor™ 488 goat anti-rabbit IgG (H+L) | Invitrogen | A11034 | 1:1000 |
| Alexa fluor™ 555 goat anti-mouse IgG (H+L) | Invitrogen | A-21429 | 1:1000 |
| Alexa fluor™ 633 goat anti-rabbit IgG (H+L) | Invitrogen | A-21070 | 1:1000 |
| Alexa fluor™ 546 donkey anti-sheep IgG (H+L) | Invitrogen | A21098 | 1:1000 |
| Alexa fluor™ 555 goat anti-rat IgG (H+L) | Invitrogen | A-21434 | 1:1000 |

**Supplementary Table 4 Primer**

All Primer were purchased from Eurofins Genomics.

| Gene | Forward Primer | Reverse Primer |
|---|---|---|
| HPRT | TGA CCT TGA TTT ATT TTG CAT ACC | CGA GCA AGA CGT TCA GTC CT |
| SeV | GGA TCA CTA GGT GAT ATC GAG C | ACC AGA CAA GAG TTT AAG AGA TAT GTA TC |
| KOS | ATG CAC CGC TAC GAC GTG AGC GC | ACC TTG ACA ATC CTG ATG TGG |
| Klf4 | TTC CTG CAT GCC AGA GGA GCC C | AAT GTA TCG AAG GTG CTC AA |
| c-Myc | TAA CTG ACT AGC AGG CTT GTC G | TCC ACA TAC AGT CCT GGA TGA TGA TG |
| TBXT | TGC TTC CCT GAG ACC CAG TT | GAT CAC TTC TTT CCT TTG CAT CAA G |
| HAND1 | TTC AAG GCT GAA CTC AAG AAG G | CTT TAA TCC TCT TCT CGA CTG GG |
| AFP | AGA AAT ACA TCC AGG AGA GCC A | TTT GTG TAA GCA ACG AGA AAC G |
| SOX17 | ATT TTG TCT GCC ACT TGA ACA G | GAA GCT GTT TTG GGA CAC ATT C |
| Tuj1 | CTC AGG GGC CTT TGG ACA TC | CAG GCA GTC GCA GTT TTC AC |
| SOX1 | CAA TGC GGG GAG GAG AAG TC | CTC GAA ACA TTT TGG GTG GGG |
| SOX10 | ATG TCA GAT GGG AAC CCA GA | GTC TTT GGG GTG GTT GGA G |
| hPAX6 | CAG ACA CAG CCC TCA CAA AC | TCA TAA CTC CGC CCA TTC AC |
| hOct4 | GGG GGT TCT ATT TGG GAA GGT A | ACC CAC TTC TGC AGC AAG GG |
| hGAPDH | AGC CAC ATC GCT CAG ACA C | GCC CAA TAC GAC CAA ATC C |
| HPRT | TGA CCT TGA TTT ATT TTG CAT ACC | CGA GCA AGA CGT TCA GTC CT |
